# Melt electrowritten scaffold architectures to mimic vasculature mechanics and control neo-tissue orientation

**DOI:** 10.1101/2023.04.17.537192

**Authors:** Angelica S. Federici, Brooke Tornifoglio, Caitríona Lally, Orquidea Garcia, Daniel J. Kelly, David A. Hoey

## Abstract

Cardiovascular disease is one of the leading causes of death worldwide, commonly associated with the development of an arteriosclerotic plaque and impairment of blood flow in arteries. Current adopted grafts to bypass the stenosed vessel fail to recapitulate the unique mechanical behaviour of native vessels, particularly in the case of small diameter vessels (<6 mm), leading to graft failure. Therefore, in this study, melt-electrowriting (MEW) was adopted to produce a range of fibrous grafts to mimic the extracellular matrix (ECM) architecture of the tunica media of vessels, in an attempt to match the mechanical and biological behaviour of the native tissue. Initially, the range of collagen architectures within the native vessel was determined, and subsequently replicated using MEW (winding angles (WA) 45°, 26.5°, 18.4°, 11.3°). These scaffolds recapitulated the anisotropic, non-linear mechanical behaviour of native carotid blood vessels. Moreover, these grafts facilitated human mesenchymal stromal/stem cell (hMSC) infiltration, differentiation, and ECM deposition that was independent of WA. The bioinspired MEW fibre architecture promoted cell alignment and preferential neo-tissue orientation in a manner similar to that seen in native tissue, particularly for WA 18.4° and 11.3°, which is a mandatory requirement for long-term survival of the regenerated tissue post-scaffold degradation. Lastly, the WA 18.4° was translated to a tubular graft and was shown to mirror the mechanical behaviour of small diameter vessels within physiological strain. Taken together, this study demonstrates the capacity to use MEW to fabricate bioinspired grafts to mimic the tunica media of vessels and recapitulate vascular mechanics which could act as a framework for small diameter graft development and functional long-term tissue regeneration.

## 1 Statement of significance

Synthetic vascular grafts utilised to bypass a blocked vessel fail to recapitulate the unique mechanical behaviour of native vessels leading to graft failure. Therefore, in this study, we determined and replicated the architecture of the vessel wall using melt electrowriting. Utilising this advanced biofabrication technology, we were able to demonstrate that these bioinspired scaffolds could recapitulate the anisotropic, non-linear mechanical behaviour of native carotid blood vessels and demonstrated that these bioinspired architectures promoted cell alignment and preferential neo-tissue orientation in a manner similar to that seen in native tissue. This approach potentially overcomes the limitations of current synthetic vascular grafts and provides a framework for vascular graft development and functional vascular tissue regeneration.

## 2 Introduction

Cardiovascular disease is one of the leading causes of mortality worldwide [1]. Most commonly the formation of an arteriosclerotic plaque leads to the obstruction of the vessel’s lumen: the arterial wall becomes stiffer, compliance is reduced, and the constriction of blood flow decreases the oxygen levels leading to tissue necrosis and ultimately patient death [2]. Treatment to restore blood flow most commonly involves the use of minimally invasive procedures such as balloon angioplasty and stenting, but if they fail, a bypass is required to divert the flow around the blockage [3]. The current *gold standard* treatment in bypass surgery remains autograft implantation [4][5]. In addition to damaging a new region of the human body, this process is also limited by graft tissue availability, restenosis, and a mechanical mismatch between venous and arterial tissue [6][7][8][9][10]. Synthetic vascular grafts (SVGs) are the most studied alternatives to autografts and demonstrate high patency for large to medium diameter vessel substitution [4][11]. However, SVGs have been shown to fail when used to replace small diameter vessels (<6 mm) [4][11], largely due to poor mechanical behaviour in terms of compliance and an inability to regenerate a homogenous endothelium, thus leading to clot formation and intimal hyperplasia [6]. Therefore, to address this unmet clinical need, it is believed that a vascular substitute that possesses properties similar to that of native vessels is essential.

A more recent alternative to SVGs are tissue engineered vascular grafts (TEVG), where a scaffold is utilised to instruct cell behaviour to drive tissue formation and/or regeneration [12]. The requirements for TEVG are numerous, but adequate compliance to match the mechanical behaviour of native tissues has emerged as a key factor in graft success [5][13][14]. Blood vessels are highly organised structures, with each component playing a fundamental role in maintaining the homeostasis of the tissue. The vessel wall can be divided into three layers; tunica intima, tunica media, and tunica adventitia. In the range of physiological pressures (80-120 mmHg), the combination of collagen and elastin fibres of the media work together to provide strength and resilience to the vessel wall and assure shape recovery during diastole [10]. This behaviour is achieved not only by the concentration of these components but importantly by their arrangement [15], where the resistance to load is given by the circumferential alignment of collagen and elastin fibres. While the specific orientation of the ECM across the vessel wall is challenging to determine due to vessel and species variability, it has been shown to decrease in alignment moving from the intima to adventitia layers (18.8° - 37.8° - 58.9° degrees) [16]. In order to ensure a TEVG possesses similar mechanical properties to the native small diameter vessel, the capacity to establish these fundamental structural aspects within a regenerative solution is a mandatory requirement for long term patency and graft success [17].

Scaffolds for vascular tissue engineering have commonly been fabricated via electrospinning, a technique that enables the deposition of small diameter fibres which simulate the native extracellular matrix (ECM). However, electrospinning has limited control over fibre architecture and is thus poorly equipped to create the highly organised structure of native vessels [4][18]. Melt electrowriting (MEW) is a recently developed additive manufacturing technique that allows for the accurate layer-by-layer deposition of fibres of micron-scale diameter, facilitating the fabrication of a highly controlled fibrous architecture in three dimensions, ultimately overcoming the limitations of classic electrospun grafts [19]. These highly organised architectures can be tuned to mimic the ECM fibre orientations found in the vessel wall [5], potentially recapitulating the mechanics of native tissues. Moreover, these MEW architectures can also act as instructional cues for infiltrating cells, where local fibres provide contact guidance controlling cell shape and neo-tissue orientation [20][21][22]. Given the versatility of MEW, this approach can be used to fabricate a wide range of geometrical cues resembling tissue specific architectures [22][23][24][25]. Indeed, within the cardiovascular space, MEW has been utilised to recreate the ECM structure of a heart valve [22] and tubular scaffolds have been fabricated to simulate a patient specific aortic root [24] and heart valve complexes [23]. Furthermore, tubular MEW scaffolds have also been fabricated, demonstrating printability up to 300 stacked layers at a winding angle (WA) of 60° [26], while others having printed auxetic tubular scaffolds and examined the mechanical behaviour of these unique architectures [27]. Very few studies have investigated these tubular scaffolds in the context of tissue regeneration. Utilising a WA of 30° to replicate a kidney tubule; van Genderen et al. demonstrated that these structures could influence cell orientation, although the ability of such scaffolds to orientate ECM was not explored [28]. Moreover, Jungst et al. demonstrated cell alignment and smooth muscle cell differentiation in a tubular MEW scaffold surrounding an electrospun lumen [29]. While these early studies demonstrate the exciting promise of MEW in the fabrication of tubular grafts, an in-depth understanding of tubular MEW architectures on the mechanical and biological performance of the VG is lacking.

Therefore, in this study we first evaluated the architecture of the native small diameter vessel wall via diffusion tensor imaging-derived tractography to establish a design criterion for fibrous scaffold architectures for TEVG. We next optimised the MEW printing parameters to generate a range of fibrous scaffolds with a structure that mimics the range of ECM fibre orientations found within the tunica media of the vessel wall. Fibrous architectures were visually and mechanically characterised, and a long-term biological study undertaken to investigate how hMSC behaviour (cell viability, orientation) and ECM deposition and organisation was impacted by scaffold architecture. Overcoming the limitations of previously reported SVGs, MEW could produce a range of scaffolds that mirrored the ECM architecture and mechanical properties of native tissue. Moreover, these ECM mimetic architectures were shown to promote hMSC infiltration and smooth muscle differentiation, leading to the deposition of neo-tissue that preferentially oriented along the long axis of the scaffold pore architecture. This robust cell infiltration, ECM deposition and vascular mimetic orientation illustrates the significant potential of these scaffold architectures for vascular tissue engineering.

## 3 Materials and methods

### 3.1 Native tissue characterisation

Porcine carotids (up to 3 months old) were collected immediately following sacrifice from a local abattoir and transported on ice to the laboratory (N=3). The isolation of the vessels was completed following the aortic pathway from the left ventricle. Once obtained, the carotid was divided in two parts proximal to the bifurcation so that half was used for histological analysis and the other half for mechanical characterisation (3.3.4). The samples were further dissected in the longitudinal and circumferential directions and fixed in 4% paraformaldehyde (PFA) overnight at 4 °C. The sections, 5 µm in thickness, were stained with H&E to assess the overall tissue organisation and cell distribution; picrosirius red staining was used for collagen identification and polarised light for collagen distribution analysis, and Verhoef-van Gieson (adapted from [30]) was used for elastin characterisation.

The investigation of microstructural alignment in native arterial tissue was performed via diffusion tensor imaging-derived tractography based on a previous optimised protocol [31]. Briefly, porcine carotid arteries of 6-month-old healthy Large White pigs were dissected, cleaned of connective tissue, then cryopreserved at a controlled rate of −1 °C/min to −80° in a tissue freezing medium. After thawing, both fresh and fixed vessels were imaged in fresh phosphate buffered saline at room temperature, with the longitudinal axis aligned parallel with the main magnetic field. Imaging, sequence, and tractography information can be found in Supplementary materials. An open-source software, ExploreDTI [32], was used for tractography visualisation. The helical angles were calculated by taking the arctan of the z-component of the first eigenvector divided by the x-component of the first eigenvector. This yields the angle between B_0_ (longitudinal axis of vessel) and the plane normal to B_0_ (the transverse plane).

### 3.2 Melt electro-writing (MEW) printer

A custom-made MEW printer was used to direct the deposition of polymeric micron-scale fibres in a preferential direction over a metal plate collector or rotating mandrel [33]. Briefly, a constant air pressure was applied to a luer-lock syringe (Nordson 3cc) so that melted PCL was extruded through a needle of 22G (Nordson 1/4"). Two thermocouples were used to control and adjust the heat for both the barrel and the needle so that a consistent and stable jet was obtained. At this point, an electric field was applied using a high voltage power supply (Heinzinger 30,000 V LNC-Series) to the needle and PCL fibres collected onto a grounded collector. Motion Perfect v4.4 was used to control all the 4-axis stepper motors motion (Arcus Technology) as well as pressure, voltage, and temperature settings. This custom setup enabled the use of a single code to determine all the variables of the process to work in synchrony and to execute the code for the different geometries.

A range of printing parameters were tested to achieve the best accuracy in fibre deposition. Pressure, temperature, voltage, and distance between spinneret and collector were varied until accuracy in the deposition, reproducibility of the scaffolds, and a 10 µm target fibre diameter was achieved.

### 3.3 Scaffold fabrication and characterisation

#### 3.3.1 Planar scaffold fabrication

With regards the MEW onto a planar collector, a melting temperature of 85 °C, pressure of 0.5 Bar, 6 kV and a distance of 5 mm between the needle and the collector were used to obtain aspect ratios of 1:1, 1:2, 1:3, 1:5 which correspond to WAs of 45°, 26.5°, 18.4°, 11.3° [34]. The space between fibres was calculated to match the targeted angle as well as to maintain a pore area comparable to the 400 µm box spacing (∼ 0.16 mm^2^). The pattern was replicated for a minimum of 30 times to achieve a scaffold of approximately 600 µm in height. In order to obtain a rhombus shape pore pattern, codes were written with diagonal translation, where both the x- and y- direction motors were moving simultaneously (**Figure 1**-C).

**Figure 1:**
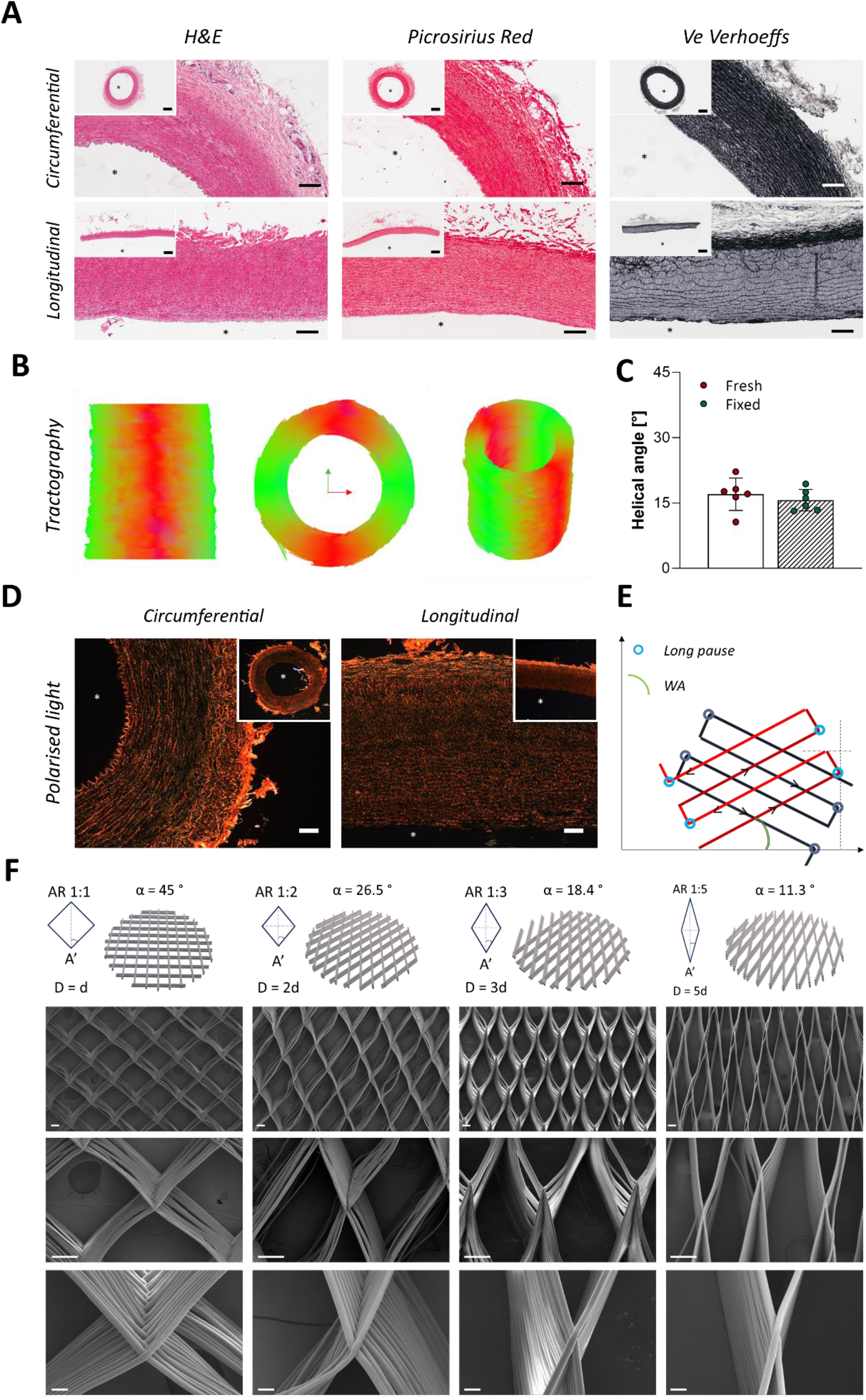
*Analysis of native porcine carotid organization to identify the ECM patterns that will be replicated in MEW scaffold architecture. **(A)** Histological sections of native samples in both circumferential and longitudinal views. Original magnification 1x and 5x. Scale bars 1 mm and 200 µm respectively. **(B)** Diffusion tensor imaging-derived tractography, which follows the dominant water diffusion direction in the tissue. Circumferential alignment is visualised by the red-green tracts, whereas axial alignment would be blue. **(C)** Quantification of the degree of alignment of the microstructure in the arterial tissue calculated based on tractography data. **(D)** Polarised light images were taken at 2x and 10x original magnification. Scale bars 200 µm. Asterisks identify the internal lumen. **(E)** Schematic representation of printing pattern where the second layer is deposited (blue) in the opposite direction of the previous one (red). **(F)** SEM images of different MEW scaffold geometries. From top to bottom rows schematic representation of desired geometries, original magnification 45x tilted at 30 degrees, 150x tilted at 30 degree and 450x straight. Scale bar respectively 10, 100 and 20 µm. From left to right 45° (AR 1:1), 26.5° (AR 1:2), 18.4° (AR 1:3), and 11.3° (AR 1:5). D and d identifying the major and minor diagonal of the rhombus geometries.*

#### 3.3.2 Tubular scaffold fabrication

With regards the MEW onto a tubular mandrel, a pressure of 0.5 Bar, 6.5 kV and total translational speed of 6 mm/h were used to obtain an aspect ratio of 1:3 that corresponds to a WA of 18.4°. A correlation between the rotational velocity and the translational speed was optimised so that an accurate deposition of 30 layers was possible. The pore area was maintained constant as per previous planar samples, ∼0.16 mm^2^.

#### 3.3.3 Planar and tubular scaffolds morphological characterisation

After a first assessment with a bright field microscope (Olympus IX83), scanning electron microscopy (SEM) was used to investigate morphological characteristics of the fibrous scaffolds. Briefly the samples were sputter coated with gold/palladium for 60 s at a current of 40 mA using a Cressington 208HR sputter coater. The scaffolds were imaged in a Zeiss ULTRA plus SEM with an accelerating voltage of 5 kV, straight and tilted at 30° at different magnifications of 45x, 150x and 450x. WA estimation was performed within the optical microscopy pictures, while fibre diameters were measured with SEM images so that at least 5 frames were analysed with 15 different measurements.

#### 3.3.4 Mechanical characterisation

A direct comparison of the mechanical behaviour was performed between native tissue and all fabricated scaffolds introduced above (3.3.1). All specimens were cut into a rectangular shape 3 mm wide and approximately 20 mm long and were then subjected to a tensile displacement at a rate of 10 mm/min until failure in dry conditions at room temperature. For vascular tissues, a supplementary conditioning phase of 10 cycles up to 5% strain with a speed of 10 mm/min was added prior to the initiation of the test. To assure the right positioning between the grips, native and synthetic samples were pre-loaded to 0.05 N and 0.01 N, respectively. For each geometry, four scaffolds were tested along both the short (longitudinal direction) and the long diagonal (circumferential direction). For biological tissues, six samples from three porcine carotids were obtained and were tested in both the longitudinal and circumferential directions. Information was extrapolated from the data obtained from the stress-strain curves such as ultimate tensile stress (UTS), young’s modulus, yield stress and yield strain, strain extension of the toe region and modulus of the toe region (**Figure S1**). The toe region was identified as the initial elastin dominated region of the typical J-shaped curve, and the young’s modulus as the secondary linear region of the curve where physiologically collagen and elastin fibres are simultaneously loaded.

A preliminary simplification was used to approximate the compliance of the scaffold, with the assumption that the change in the circumference is proportional to the change in diameter, it was possible to apply Laplace’s law as follows:

*Equation 1*

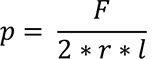

Where p is the luminal pressure, *F* is the loading force, *r* is the radius, and *l* is the width of the samples used. Laplace’s law was used to establish the force at 80 and 120 mmHg when the radius corresponds to the unload sample – grip to grip separation. At this point the resultant travelling distance obtained during the test at that specific force value could be detected and therefore the diametrical change obtained. Values of estimated compliance were then calculated with [35]:

*Equation 2*

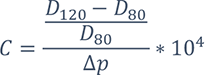

Where D corresponds to the diameters respectively at 120 mmHg and 80 mmHg, and Δp is the pressure differential (i.e. 40 mmHg).

A second mechanical test to assess the dynamic response of the scaffolds was also performed [22]. Briefly, a pre-load of 0.01 N was used to assure the right positioning of the specimens; then 12 cycles with a 5% strain increase up to a maximum 60% strain at a constant velocity of 10 mm/min and a load removal down to 0% was applied. The area of hysteresis was calculated through a custom-made MATLAB® script that allows the calculation of the ratio between the areas of unloading and loading of the samples [36]. As the areas under the curves are representative of the elasticity of the samples, a decreasing ratio is evidence of an increasing hysteresis, meaning the elasticity of the construct is lost [22].

Mechanical characterisation of tubular scaffolds was performed through the ring test method. Briefly, cylindrical specimens were cut and dimensions recorded for further analysis. A pre-load of 0.01 N was applied for a correct positioning before starting the test. A 200 N load cell was used to record the force response of the samples undergoing a stretch until failure at a constant velocity of 10 mm/min. This procedure was performed for both synthetic and native tissue samples. In the case of native porcine carotid specimens a pre-conditioning up to 5% strain was applied 10 times so that a correct rearrangement of collagen fibres was completed before the actual test. The same information reported above were extrapolated from the curves and compared.

### 3.4 Cell proliferation and extra cellular matrix (ECM) deposition

#### 3.4.1 hMSC cell culture

Human mesenchymal stem/stromal cells (hMSC) were harvested from a sacral second anterior iliac (S2AI) following ethical approval from the Institutional Review Board (IRB) of the Mater Misericordiae University Hospital (IRB no. 1/378/2038). HMSCs were cultured in low glucose Dulbecco’s modified eagle’s medium (DMEM, Sigma) supplemented with 10% fetal bovine serum (FBS) (Biosera lot#11307) and 5% penicillin and streptomycin (Sigma Aldrich). HMSCs were expanded in T175 flask at 37 °C, 95% humidity and 5% CO_2_ and passaged when 80% confluent.

#### 3.4.2 Evaluation of seeding technique

Samples were treated with a 2 M NaOH solution for 1 hour to increase hydrophilicity of the PCL construct [37]; they were then sterilised with UV for 20 minutes on each side, and then hydrated with a decreasing concentration of ethanol, specifically 100%, 90%, and 70% for 15 minutes each step. Finally, specimens were soaked in fresh growth media consisting of DMEM with 10% FBS overnight for preconditioning. A pilot biological study was performed to assess the confinement seeding efficacy of the PDMS mould over the scaffolds, two chemical treatments to enhance PCL hydrophilicity, and to evaluate the difference between two seeding densities - 1*10^5^ and 5*10^5^ cells/scaffold.

#### 3.4.3 Evaluation of cell viability

Live/dead analysis was performed; briefly calcein AM 4 mM staining (Cambridge Biosciences BT80011-1) and ethidium homodimer 2 mM (Cambridge Biosciences BT40014) were used in a concentration respectively of 10 µl and 2.5 µl over 5 ml of PBS. Fluorescent imaging was carried out using a Leica SP8 scanning confocal microscope at different magnifications.

#### 3.4.4 Evaluation of tissue maturation

Samples were placed inside a custom-made PDMS mould to confine the seeding volume for the first 24 hours and seeded with hMSCs at 5*10^5^ cells/scaffold. Scaffolds were then moved into fresh well-plates and media enriched with 10 ng/ml TGFβ1 to induce SMC differentiation following a previously optimised protocol (3.4.2). Every 7 days samples were imaged for live dead analysis. Samples were collected at days 1, 14, and 28 for immunohistochemical characterisation as well as histological analysis to assess cell distribution and differentiation throughout the scaffolds. Moreover, cell number and ECM deposition was evaluated through biochemical analysis. The results were compared to non-TGFβ1 treated controls.

#### 3.4.5 Immunochemical staining and analysis

Briefly, cells were permeabilised in a 0.1% (v/v) solution of Triton X-100 for 10 minutes followed by a wash in PBS and 1 hour incubation in 1% (w/v) bovine serum albumin (BSA) solution in PBS. Primary antibodies were left in incubation overnight at 4 °C, followed by secondary antibodies incubation for 1 hour in the dark. Phalloidin (Biosciences Alexa 488, A12379) and DAPI (Sigma 32670-5MG-F) staining was completed for 20 minutes and 5 minutes respectively at a concentration of 1:40 and 1:2000.

Samples were stained with two different primary antibodies chosen as markers of the different stages of vascular smooth muscle cell differentiation: αSMA and calponin (**Table 1**). Vinculin was also adopted as a marker of focal adhesion assembly.

**Table 1:** *Summary of all antibodies used and their relative work’s concentrations.*

| Antibody | Supplier | Reference | Dilution |
| --- | --- | --- | --- |
| αSMA | Abcam | ab21027 | 1:200 |
| Calponin | Abcam | ab700 | 1:300 |
| Anti-mouse IgG Alexa Fluor 594 | Life Technologies | A21203 | 1:500 |
| AlexaFluor488-Phalloidin | Life Technologies | A12379 | 1:40 |

Fluorescent imaging was carried out using an Olympus IX83 epi fluorescent microscope and a Leica SP8 scanning confocal microscope. The intensities of the antibodies were calculated with a custom-made MATLAB® script. Briefly, a constant threshold was imposed to all the images; multiple ROI were identified as the areas where f-actin staining was expressed, and the intensity measured within the ROI while the outside region was considered as background intensity. The obtained results were normalised over the number of cells stained with DAPI. The procedure was optimised based on a previous study [38].

#### 3.4.6 Biochemical analysis

For DNA, collagen, and glycosaminoglycans (GAG) content, scaffolds were enzymatically digested in 400 µl of papain solution. Samples were then incubated at 60 °C overnight under constant rotation. DNA content was quantified using a Quant-iT™ Pico Green™ dsDNA Assay Kit (Invitrogen, P7589), with excitation and emission wavelengths of 485 nm and 528 nm, respectively. The amount of sulphated glycosaminoglycan (sGAG) was determined using the dimethyl methylene blue dye-binding assay. Total collagen content was accessed through the quantification of hydroxyproline content. Briefly, for 18 hours constructs were hydrolysed in 38% HCl at 110 °C after which they were analysed via chloramine-T assay and quantified with a ratio of 1:7.69 hydroxyproline:collagen respectively [39].

#### 3.4.7 Histological analysis

Fixed samples were embedded in 4% agarose solution to confine the constructs while xylene was used to dissolve the PCL fibres throughout the construct. Scaffolds were then dehydrated, and paraffin embedded. Sections of 5 µm were collected and Haematoxylin and Eosin stain was used to assess overall tissue formation (H&E), Alcian blue for sGAG content and Picrosirius red for collagen content, using a Leica 5010 Auto Stainer. Images were acquired with Scan Scope (Aperio Technologies Inc., USA). Optical density of these stains was performed through a colour deconvolution plug-in in ImageJ.

#### 3.4.8 SEM biological sample preparation and analysis

Constructs were fixed in a solution of 3% glutaraldehyde (GTA) in 0.1 M cacodylate buffer at 4 °C for a minimum of 12 hours. When all the samples were collected and fixed, an incubation of 10 minutes in 0.1 M cacodylate buffer was performed twice to wash the unreacted GTA. A dehydration in ethanol series was performed from 50 to 100% with a step at 70% and one at 90%, immersion was prolonged for 10 minutes and repeated two times. Hexamethyldisilazane (HMDS) was used to cover samples for one hour with a change of solution midpoint after which the samples were mounted onto stubs so that the shrinking process was limited. Scaffolds were left to dry in a fume hood at least overnight before proceeding with the imaging with a SEM at different magnifications (3.3.3).

### 3.5 Statistical analysis

The number of scientific replicates was identified with the letter N and the technical replicates with the letter n, throughout the text. For statistical analysis GraphPad Prism 8 (GraphPad Software, USA) was used for all the data. Where possible, One-way ANOVA test was performed in combination with Dunnett’s multiple comparison. The results are presented as mean and standard deviation where appropriate.

## 4 Results

### 4.1 Vascular tissue characterisation

Initially, to develop a design criterion for a vascular graft and to build on data from the literature, a histological and mechanical characterisation of a porcine carotid artery was performed. Histological analysis demonstrated the subdivision of the vessel wall into three distinct layers (**Figure 1**). The overall thickness of the wall was approximately 770.5 ± 117 µm, with each layer from outside to inside being 117.6 ± 46 µm, 658.7 ± 126 µm, and 19.7 ± 8.2 µm, respectively. Furthermore, a significant difference in terms of lumen size was observed among the same segment of the carotid in relation to the distance from the bifurcation of the aorta pathway. The average external diameter was calculated to be 3.8 ± 1.1 mm and the internal to be 2.4 ± 1 mm. Elastin staining, (black coloured fibres between tunica media and adventitia) and collagen staining (picrosirius red coloured fibres) pointed out the evident concentration of elastic components predominately in the media layer of the vessel wall (**Figure 1**-A). To better assess collagen fibre organisation within the vessel wall, 3D visualisation of the alignment in fresh and fixed native porcine carotids was modelled by diffusion tensor imaging derived tractography (**Figure 1**-B), where circumferential alignment can be visualised by red-green tracts. There was no difference between fresh and fixed vessels. Mean helical angles confirmed this circumferential alignment, with fresh vessels having a mean helical angle of 15.7 ± 2.6° [max 18.3° – min 13.1°] and fixed 17± 2.6° [max 19.6° – min 14.4°] (**Figure 1**-C). Polarised light further clarified the specific organisation of the collagen fibres along the circumferential direction that provides mechanical resistance to blood pressure (**Figure 1**-D).

### 4.2 MEW scaffold fabrication and mechanical characterisation

All the targeted architectures were successfully printed with high accuracy (**Figure 1**-F). The optimal processing parameters of the MEW printer were identified to reliably fabricate scaffolds with a WA of 45° (AR 1:1), 26.5° (AR 1:2), 18° (AR 1:3), and 11° (AR 1:5), which covers the range of collagen architectures (helical angles) identified in **Figure 1**-B, C. An average fibre diameter of 8.9 ± 0.5 µm (N=4, n=20) was observed for all geometries via SEM images (**Figure 1**-F) and the height of the scaffold walls (30-layers) was approximately 550 µm, which rose closer to the theoretical target of 600 µm at the crossing points, and is similar in height to the width of the tunica media identified in Figure 1. The resulting fibres were found to be well stacked one over the other with well-defined crossing points. Analysis of the printed scaffolds demonstrated that angles of 53.4 ± 2.6°, 38.5 ± 3.5°, 23.4 ± 2.8° were obtained for WA 26.5°, WA 18.4°, and WA 11.3° respectively, demonstrating the accuracy of this printing approach. The scaffold with a WA 45° was perfectly fabricated given the simplicity of the architecture.

#### 4.2.1 Mechanical characterisation

The different fibre architectures fabricated via MEW demonstrated unique mechanical behaviour. Uniaxial tensile testing was performed on strips of each scaffold in both ‘circumferential (long pore axis)’ and ‘longitudinal (short pore axis)’ directions (**Figure 2**-A-B) and compared to native tissue controls. PCL alone does not possess the anisotropic mechanical behaviour that would mimic that of a native vessel [40][41], however by melt electrowriting fibrous architectures an anisotropic behaviour could be obtained, with increased anisotropy observed as the angle between the fibres decreases, leading to a more physiological response (**Figure 2**-C-i-iv). The mechanical properties of all scaffold designs are reported in **Table S2** and represented in **Figure S2** for both directions of the applied force, as well as for the native tissue.

**Figure 2:**
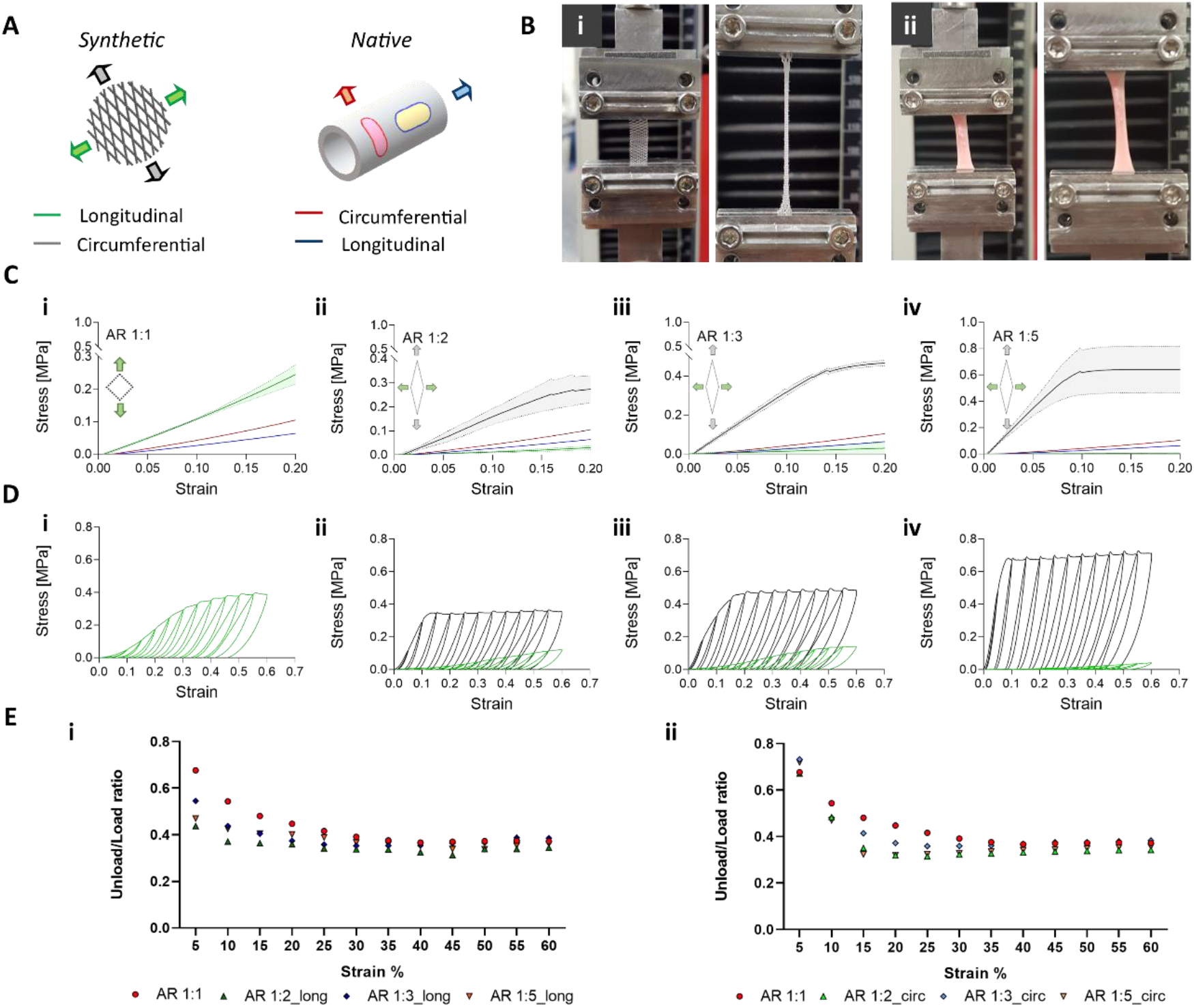
*Mechanical characterisation of MEW synthetic grafts and comparison with native vascular tissue through uniaxial tensile testing. **(A)** Schematic representation of longitudinal and circumferential definition of scaffolds orientation. **(B)** Representative images of tested strips for both **(i)** synthetic and **(ii)** native samples before and during the test. **(C)** Stress strain curves in both longitudinal and circumferential direction of the different geometries compared to native tissue response. **(i)** AR 1:1, **(ii)** AR 1:2, **(iii)** AR 1:3, **(iv)** AR 1:5. Data show Mean ± SD (N=4, n= 1). **(D)** Representative stress strain curves for all the geometries under cyclic incremental loading from 5% to 60% strain in both longitudinal and circumferential direction. **(i)** AR 1:1, **(ii)** AR 1:2, **(iii)** AR 1:3, **(iv)** AR 1:5. **(E)** Load-Unload ratios for all the geometries in both **(i)** longitudinal and **(ii)** circumferential directions. Data presented just with mean value (N=4).*

In the longitudinal direction, the linear region (young’s modulus) was comparable for all samples while the UTS decreased with a reduction in the WA (**Figure 2**-C-i-iv and **Figure S2**-A-i-ii). On the contrary, in the circumferential direction, a reduction in the WA resulted in an increase in both the young’s modulus and the UTS (**Figure 2**-C-i-iv and **Figure S2**-A-i-ii). When force was applied in the longitudinal direction decreasing the WA from 45° to 11° degrees increased the extension of the elastin dominant region (toe region) (**Figure S2**-A-iii). The yield strain followed the same increasing trend when the WA was reduced in the longitudinal direction (**Figure 2**-C-i-iv and **Figure S2**-A-v). Oppositely, in the circumferential direction the WA did not drastically affect the elastin dominant region (toe region), with all values below 15% (**Figure 2**-C-i-iv and **Figure S2**-A-iii). In this case, the yield strain obtained decreased with a decreasing WA between fibres (**Figure 2**-C-i-iv and **Figure S2**- A-v).

The compliance (defined as the capacity of samples to recover after deformation) of the MEW structures was also assessed in the longitudinal and circumferential directions (**Table S2**). In the longitudinal direction, the compliance approximation was not detected for the diamond-shape architectures. In contrast, in the circumferential direction, reducing the WA resulted in a reduction of the compliance values from 5 to 2 (1/100) mmHg.

When comparing the obtained results with native tissue data, the rhombus architectures were able to generate stress-strain curves that resembled the typical J-shape curve of the native tissue within the range of 0-20% strain (**Figure 2-**C-i-iv). In the longitudinal direction, values in the secondary linear region were lower, but no significant difference was detected. Reducing the WA in the longitudinal direction, resulted in a significant decrease of UTS values compared to native carotid (**Figure S2**-A-i). In the circumferential direction reducing the WA led to a 5-fold increase of young’s modulus (relative to the WA-45°), while the UTS values remained in the range of native tissue with exception of WA-26° (**Figure S2**-A-i). A closer resemblance of native tissue mechanical properties is observed in the longitudinal direction compared to the circumferential direction. The WA-18° scaffold design, which closely mirrors the collagen architecture within the tunica media (**Figure 1**- C– 15.7°), better matches native tissue mechanical behaviour for physiological levels of strain.

We next looked to investigate the ability of our grafts to withstand repeated loadings as would occur *in vivo* (**Figure 2**-D-i-iv). The response of the MEW scaffolds to cyclic loading (12 cycles from 5% up to 60% strain at a constant velocity of 10 mm/min) was different when the samples were loaded in circumferential and longitudinal directions (**Figure 2**-D). In the circumferential direction, reducing the WA was found to result in higher stress values for a given level of applied strain and the plateau was reached at lower strain values. On the contrary, an overall lower level of hysteresis was observed for scaffolds tested in the longitudinal direction, with values of stress below 0.1 MPa reached for smaller Was identifying a weaker resistance (**Figure 2**-D). The ratio between the area of unloading and loading was obtained with a custom-made MATLAB® script and the data showed an early occurrence of plastic deformation for all samples (**Figure 2**-E). In the longitudinal direction (**Figure 2**-E-i), all the geometries underwent plastic deformation before 15% strain. On the other hand, for circumferentially loaded specimens (**Figure 2**-E-ii) decreasing the WA led to an earlier occurrence of plastic deformation at almost 10% of the strain.

To summarise, the data reported demonstrates that the investigated architectures tune anisotropic mechanical behaviour with an increased divergence between longitudinal and circumferential data with a reduction in the WA. Of all architectures investigated, the WA-18° best replicates the mechanical performance of native vessels.

### 4.3 Cell behaviour and extra cellular matrix (ECM) deposition

#### 4.3.1 Cell viability and differentiation

The pilot evaluation of cell seeding methods identified that a higher seeding density (5*10^5^ cells/scaffold) in a confined custom made PDMS mould was the most effective approach, where hydrophilic samples demonstrated good cell viability and infiltration (**Figure S3**). Utilising this higher seeding density, hMSC proliferation, differentiation, alignment, and ECM deposition and orientation were assessed over a 28-day study. The presence of TGFβ1 did not alter the adhesion capacity of the cells to the scaffold compared to the standard growth media, as demonstrated by live/dead analysis at day 1 (**Figure 3**-A). Data demonstrated no significant difference in cell seeding efficiency among samples except for WA-11° (AR 1:5), where a lower viability was observed only at day 1 (**Figure 3**-B). The overall DNA content of the cell seeded constructs was observed to reduce with time in culture (**Figure 3**-C). Overall, no significance difference in DNA content was observed between AR 1:1, AR 1:3 and AR 1:5 at day 28. Positive staining for the vascular smooth muscle cell differentiation marker αSMA was observed at day 14 and day 28, with greater cell alignment and intensity recorded for lower ARs (**Figure 3**-D and **Figure S4**). Thus, a valid protocol for scaffold seeding and cell viability was identified, and the initiation of SMC differentiation was observed for all rhombus like architectures (**Figure 3**), although no architecture demonstrated a clear advantage in terms of SMC differentiation.

**Figure 3:**
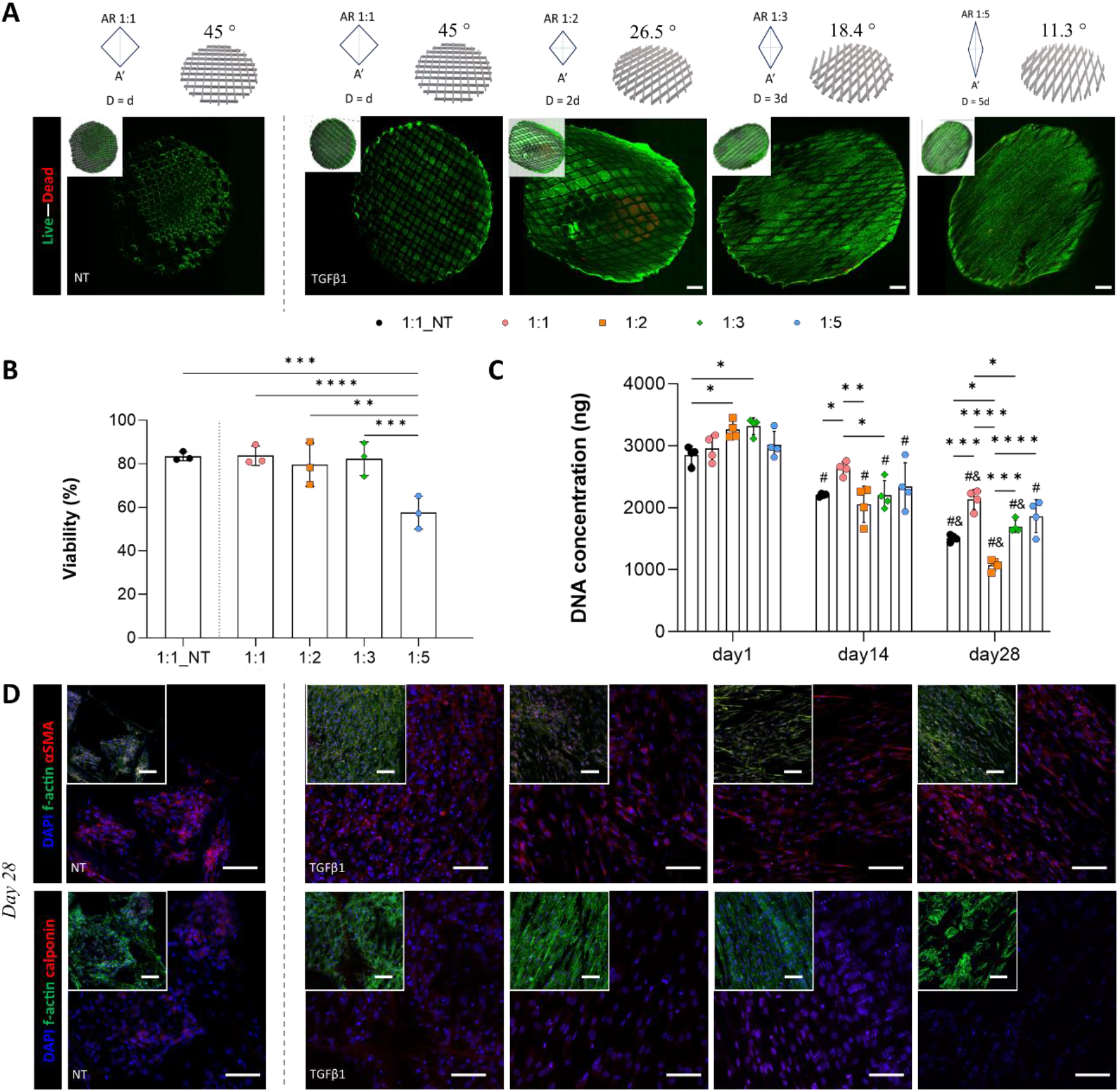
*Biological characterisation of cell behaviour in melt-electrowritten (MEW) scaffolds. **(A)** Live dead images representative of each geometry at day 28 for both non-treated samples (NT) and treated samples with TGFβ1. Scale bar 500 µm. **(B)** Viability quantification from live dead samples at day 1. **(C)** DNA content for all the groups in comparison at day 1-14-28. P value *≤ 0.05, **≤ 0.005, ***≤ 0.001, **** ≤ 0.0001. Evaluation of single group evolution at different time point identified by # statistically different from day 1, & statistically different from day 14. **(D)** IHC staining for cell differentiation over MEW samples at day 28. Scale bar 100 µm.*

#### 4.3.2 Cell orientation

Utilising F-actin as an indicator of cell orientation, the alignment of the cells along the major diagonal of the MEW scaffold frame was clearly visible, particularly with the lower WA architectures (**Figure 4**). At day 1 there was no preferential direction of cell orientation, with the majority of cells attaching to MEW fibres and not infiltrating into the centre of the pores. Over time, cells proliferated to occupy the entire pore area and began to orientate along the main axis of the pore demonstrating contact guidance (**Figure 4**-A). A further directionality analysis highlighted an increase in orientation, particularly from days 14 to 28 (**Figure 4**-B,C). Interestingly, squared geometries reported two peaks around 45° as per their WA values, thus demonstrating an orthogonally oriented pattern with no preferential orientation. On the contrary, the narrower the WA the higher was the resulting alignment with minimum dispersion recorded (**Figure 4-**D). The cellular alignment in the WA-18° (dispersion 16.2 ± 5°) and WA-11° (dispersion 14.6 ± 2.5°) scaffolds better mimicked the native vascular alignment (helical angle 15.7 ± 2.6°) (**Figure 4**).

**Figure 4:**
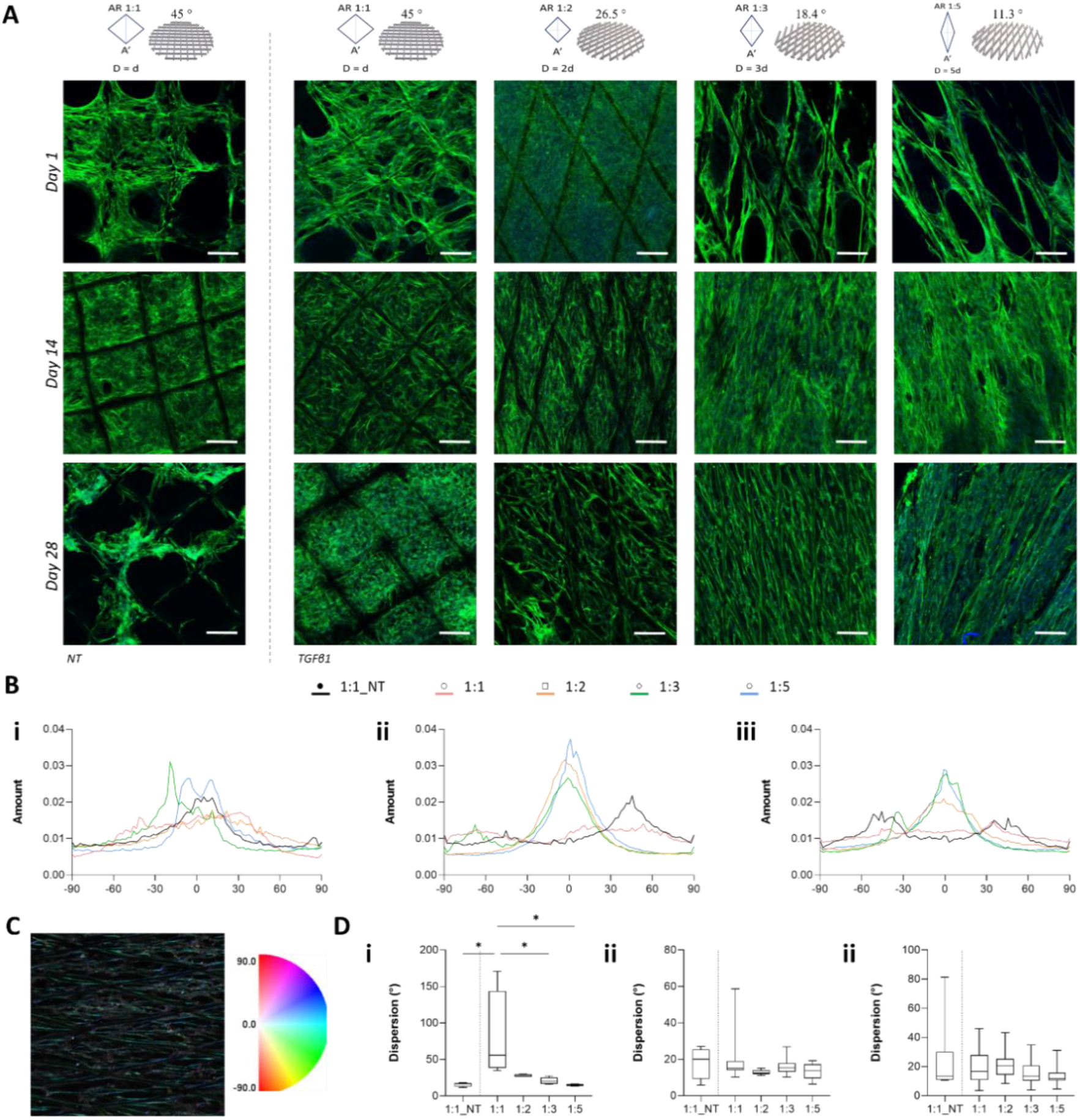
*Cell orientation analysis **(A)** Representative images of F-actin (green) and DAPI (blue) staining for all groups at different time points (day1, 14, 28). Scale bar 100 µm. **(B)** Directionality analysis of f-actin fibres at **(i)** day 1 (n=4, N=1), **(ii)** day 14 (n=4, N=3) and **(iii)** day 28 (n=4, N=3). Graph showing the amount of fibres with the same orientation and directionality preference. **(C)** Representative images of colour vector formation and associated colour wheel based on phalloidin staining orientation. **(D)** Dispersion value for all the different geometries in degrees at **(i)** day 1 (n=4, N=1), **(ii)** day 14 (n=4, N=3) and **(iii)** day 28 (n=4, N=3). P value *≤ 0.05, **≤ 0.005, ***≤ 0.001, **** ≤ 0.0001.*

#### 4.3.3 ECM deposition and orientation

ECM production was positively enhanced by TGFβ1 treatment, with treated scaffolds displaying a significant increase in collagen deposition, yet no difference was identified among the different scaffold architectures. H&E staining revealed the presence of cells throughout all scaffolds by day 28, which is consistent with the immunostaining data (**Figure 5**-A). Collagen production was highlighted by the red filaments presented in the picrosirius red stained samples (**Figure 5**-B). In detail, the collagen fibres were predominately deposited along with PCL fibres, following the underlying scaffold architecture by day 28. Collagen deposition was also quantified over time and a positive trend of increased collagen was evident with time in culture. Interestingly, while TGFβ1 treatment enhanced collagen deposition by day 14 when compared to untreated groups, there was no impact of scaffold architecture on the quantity of ECM deposition (**Figure 5**-D, E and **Figure S5**). Glycosaminoglycans (GAGs) were also detected within the constructs, as evident by Alcian Blue staining (**Figure 5**-C), however this staining was minimal and upon quantification there was no significant increases in GAGs detected over time, in response to TGFβ1, or clear trends across the different scaffold architectures (**Figure 5-**F, G). In summary, this data demonstrates that TGFβ1 is a potent stimulus to drive collagen ECM deposition in MEW scaffolds and that MEW scaffold architectures analysed here do not influence the quantity of ECM deposition.

**Figure 5:**
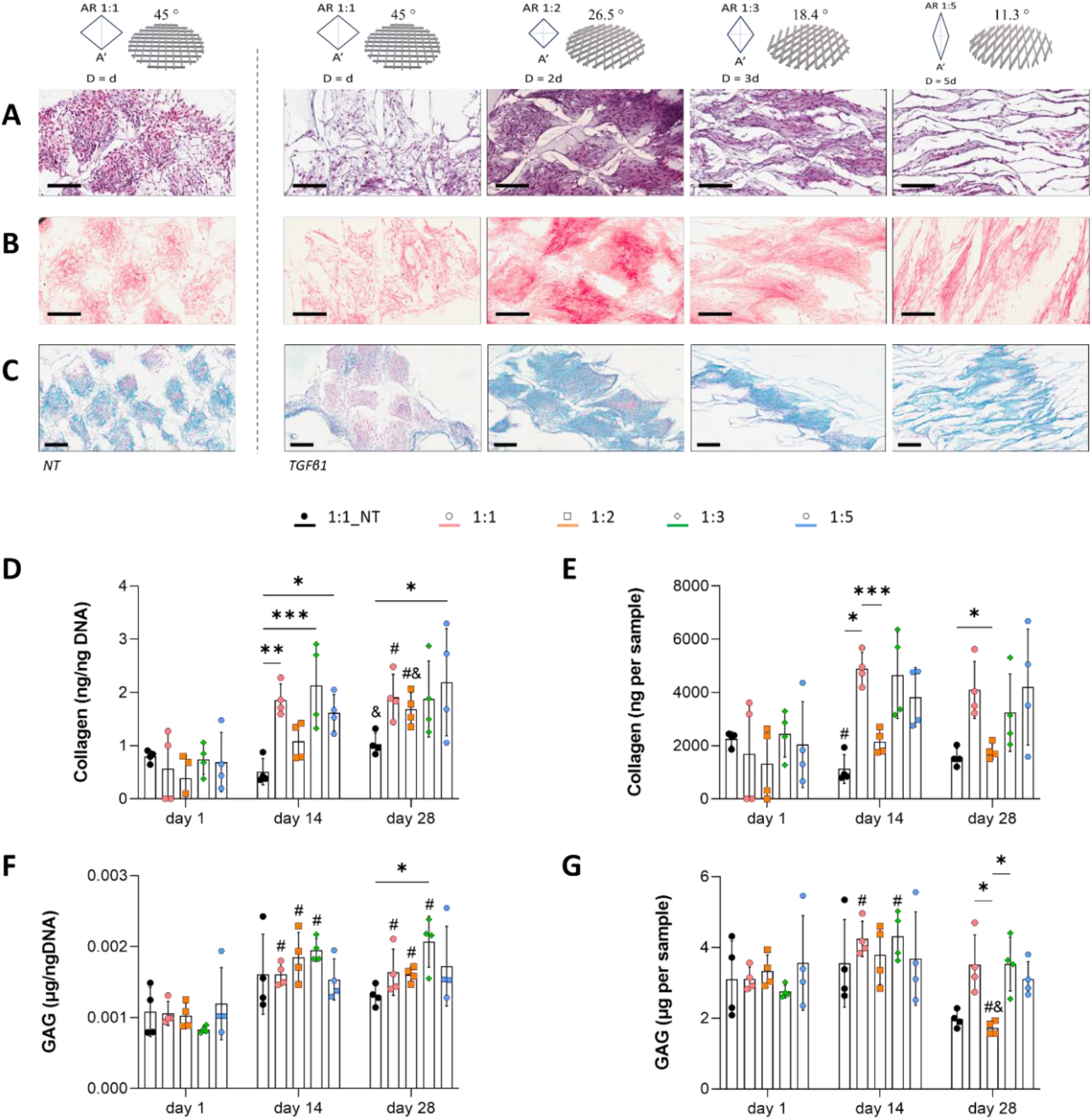
*ECM deposition analysis through histological staining of melt-electrowritten constructs at day 28 with **(A)** H&E staining, **(B)** Picrosirius Red staining, **(C)** Alcian Blue staining. Scale bar 200 µm. **(D)** Collagen content for all the groups in comparison at day 1, 14, 28 normalised for the respective content of DNA. **(E)** Collagen content for all the groups in comparison at day 1, 14, 28. **(F)** GAG content for all the groups in comparison at day 1, 14, 28 normalised for the respective content of DNA. **(G)** GAG content for all the groups in comparison at day 1, 14, 28. **(H)** Optical density value for H&E staining for all groups cultured for 14 and 28 days. **(I)** Optical density value for Picrosirius Red staining for all groups cultured for 14 and 28 days. **(L)** Optical density value for Alcian Blue staining for all groups cultured for 14 and 28 days. P value *≤ 0.05, **≤ 0.005, ***≤ 0.001, **** ≤ 0.0001. Evaluation of single group evolution at different time point identified by # statistically different from day 1, & statistically different from day 14. NT denotes not treated with TGFβ.*

While scaffold architecture did not influence the quantity of ECM produced over time, the orientation of the deposited ECM was significantly affected (**Figure 6**). SEM images after 28 days demonstrated a complete filling of the MEW scaffolds with engineered tissue for TGFβ1 treated constructs (**Figure 6**-A). Moreover, the pore architecture orientated the neo-tissue along the major diagonal, with greater matrix elongation achieved with smaller WAs. Polarised light images confirmed that not only cells but also deposited ECM followed the boundary conditions imposed by the MEW framework (**Figure 6**-B). Quantification of collagen fibre orientation demonstrated that the narrower the WA, the higher is the preferential distribution of collagen fibres along the major diagonal of the rhombus shaped samples (**Figure 6**-C). Once again, the WA-18° (dispersion 16.1 ± 9°) and WA-11° (dispersion 14.6 ± 8°) scaffolds better mimicked the native vascular collagen alignment (helical angle 15.7 ± 2.6°) (**Figure 6**). These findings demonstrate that MEW scaffolds not only influence cell alignment via contact guidance, but also neo-ECM organisation. Together, this data suggests that the WA-18° was the best performing in all aspects, enabling higher cell viability and ECM production, along with cell alignment and collagen fibre organisation which matched the circumferential orientation identified via tractography in native biological vessels (**Figure 1**-B,C) [15][42].

**Figure 6:**
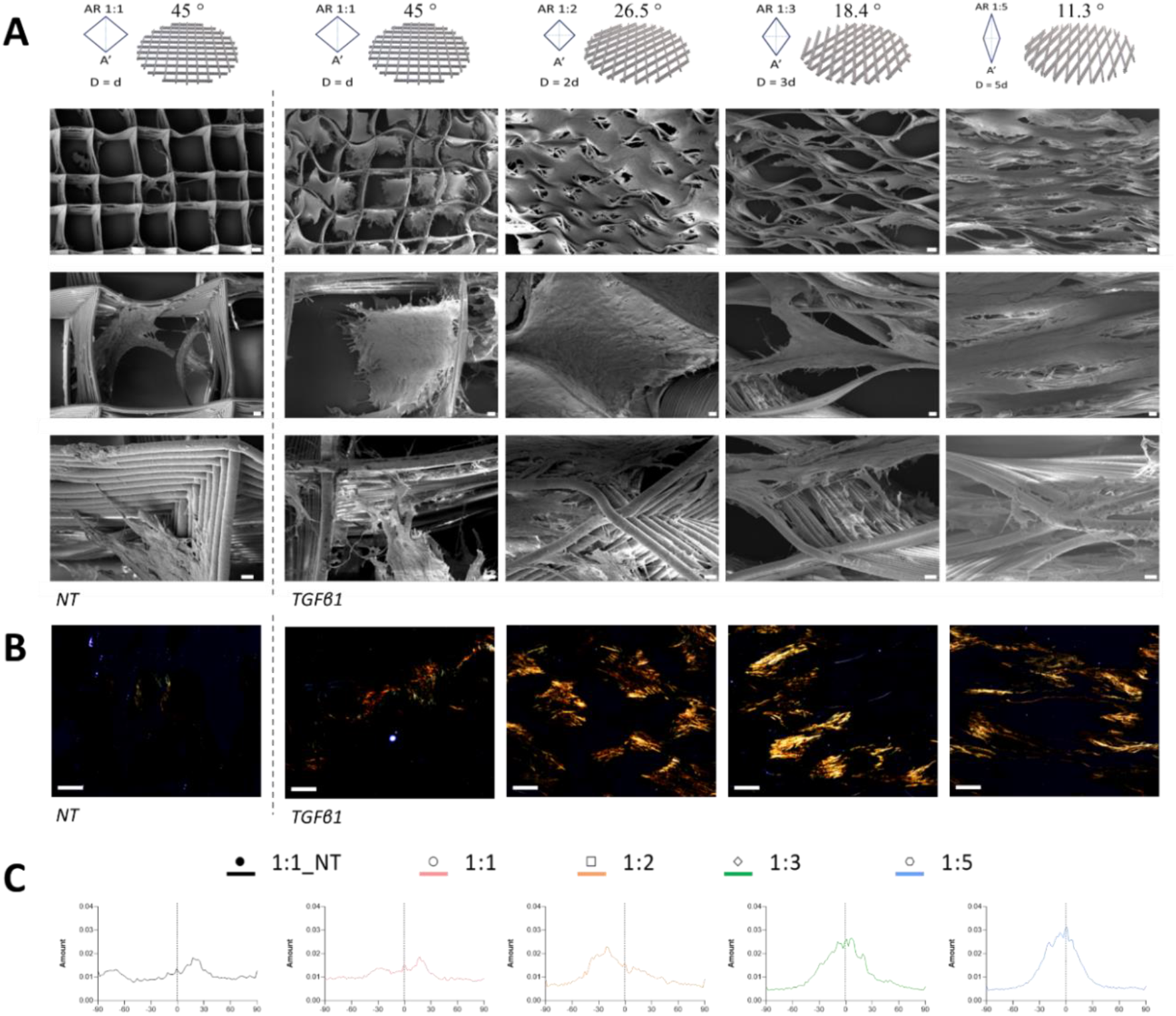
*ECM orientation analysis and quantification. **(A)** Representative SEM images of melt-electrowritten scaffold cultured for 28 days. Scale bar 100, 20, 10 µm respectively. **(B)** Polarised light images for all the groups cultured for 28 days. Scale bar 200 µm. **(C)** Directionality analysis of collagen fibres. Graph showing the quantity of fibres with the same orientation and directionality preference. NT denotes not treated with TGFβ1.*

### 4.4 Tubular scaffold fabrication and characterisation

Having identified suitable fibre architectures which recapitulate the mechanical performance of native vessels and act as boundaries for oriented neo-tissue formation, we next sought to fabricate tubular scaffolds to better represent the tunica media structure of a blood vessel. Tubular grafts with a precise and accurate deposition of PCL fibres were obtained with a WA-18° (AR 1:3), resulting in the development of scaffolds mimicking aspects of the mechanical behaviour of native vessels within the physiological range of interest (**Figure 7**). SEM images demonstrated the successful print and the accuracy in stacking layer after layer on a tubular collector matching WA-18° (**Figure 7**-A).

**Figure 7:**
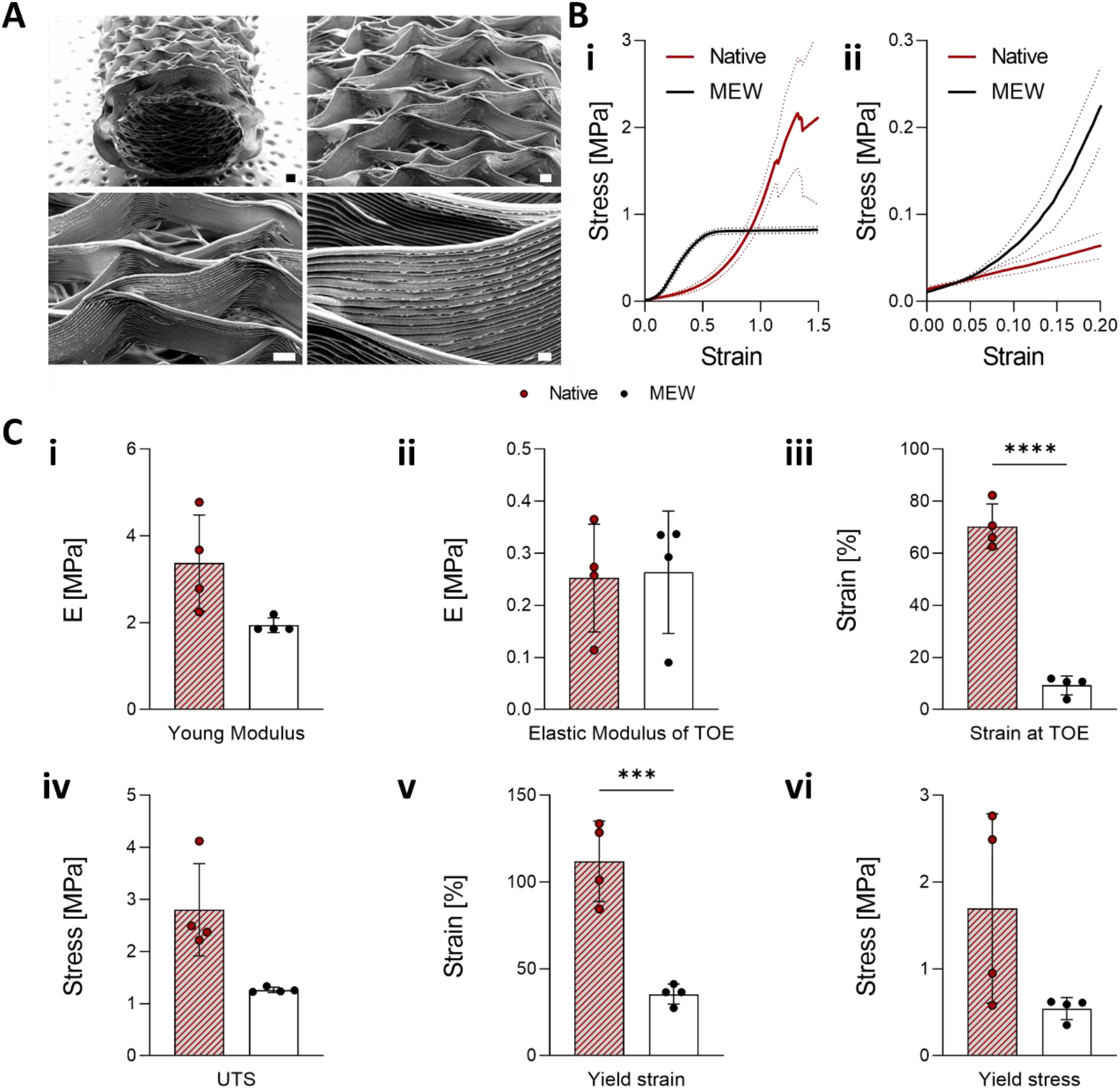
*Tubular MEW grafts with AR 1:3 matches the mechanical properties of native tissue. **(A)** SEM images representative of a tubular sample. Scale bars 200 – 100 – 100 – 20 µm respectively. **(B)** Stress-strain curves of tubular grafts of AR 1:3 in relationship to native tissue (ring test) **(i)** overall and **(ii)** zoomed in section of physiological relevant interval. **(C)** Ring test data obtained after the analysis of the stress strain curves for tubular MEW grafts and native tissue. Here reported the **(i)** young’s modulus, **(ii)** elastic modulus of the toe region, **(iii)** strain value at toe region, **(iv)** ultimate tensile strength (UTS), yield value of both **(v)** strain and **(vi)** stress. P value *≤ 0.05, **≤ 0.005, ***≤ 0.001, **** ≤ 0.0001.*

Mechanical response during a uniaxial tensile test was determined (**Figure 7**-B) and MEW scaffolds in a tubular conformation were able to match the native tissue J-shape up to 10% strain which is considered the physiological range [43], improving upon that identified in the planar conformation. Indeed, data extracted from stress-strain curves reported almost no differences between native and synthetic specimens within this range (**Figure 7**-C). Both the initial (elastic modulus of the toe region), the second linear region (young’s modulus), ultimate tensile stress (UTS), and yield stress were comparable to the native tissue (**Figure 7**-C-i-ii-iii-vi). However, strain value at toe and yield strain were significantly different from porcine carotid values (**Figure 7**-C-iv-v). To summarise, tubular grafts mimicking the native vessel tunica media architecture were fabricated and were found to closely match the mechanical properties of native vessels in the physiological relevant intervals of strain.

## 5 Discussion

Synthetic and TEVG fail to recapitulate the physiological properties of small diameter vessels resulting in failure post-implantation [18][44]. Therefore, an alternative solution is to mimic the native properties of the vessel wall to ensure long term survival and regeneration of vascular tissue.

Blood vessels are highly organised structures consisting of different layers and cell populations which provide unique mechanical properties required to maintain physiological blood pressure and flow [18][42][45]. Failing to recapitulate the compliance response of native vessels has been identified as a leading cause of graft failure. In this study we outline the initial design of a VG with a micro-architecture that mimicked the tunica media wall architecture, recapitulated native tissue mechanics, and that facilitated cell infiltration, orientation and preferential ECM deposition and organisation. Here, a range of five different PCL fibrous scaffolds were fabricated with high accuracy utilising a custom-made MEW printer [33], whereby the adopted architectures were inspired by the orientation of collagen fibres present in the tunica media of blood vessels identified using diffusion tensor imaging-derived tractography [42]. MEW scaffolds demonstrated anisotropic mechanical behaviour, which closely mimicked that of native tissue within the physiological strain range, reproducing the typical J-shape of native tissue. Samples also demonstrated resistance to cyclic loading, highlighting their potential application in continuously pressurised *in vivo* environments. Moreover, the chosen architectures allowed for abundant cell infiltration and preferential cell orientation along the major diagonal of the pore units, consistent with cellular orientation *in vivo*. Interestingly, although scaffold architecture did not influence the total amount of deposited ECM, it was evident that such deposition also followed the major diagonal of the pore units, mimicking the circumferential configuration within the tunica media, and brings confidence to the longer-term survival of the regenerated tissue post scaffold degradation. These findings were then translated to the fabrication of a tubular graft prototype, which highlighted a J-shape stress-strain curve matching native vessels properties in the elastin-dominant region. Taken together, this study demonstrates the versatility of MEW in the fabrication of scaffolds that mimics the native ECM architecture and mechanics of the vessel wall and provides guidance for cell and neo-tissue organisation.

MEW fibrous architectures were manufactured through the optimisation of printing parameters such as pressure, voltage and transitional velocity [19][46][47][48][49]. Different WAs between fibres were fabricated based on an analysis of ECM components and architecture of the native tissue wall [5][42], which have shown a degree of ECM alignment ranging from 18° to 60° degrees [18]. This was further confirmed by our findings via diffusion tensor imaging-derived tractography (helical angles of 15-17°). MEW enabled a higher accuracy in fibre deposition compared to the more commonly adopted electrospinning methods. An average fibre diameter of ∼10 µm was obtained, that is both more suitable for cell culture and more physiologically relevant than that obtained with fuse-deposition-modelling (FDM) [19][50]. Moreover, the obtained scaffold height is significantly higher than other MEW-printed scaffolds [51][52], where previous diamond shaped scaffolds with an angle of 45° degree was obtained up to 5 [53] and 10 layers [54]. The ∼600 µm of scaffold thickness achieved here matched the size of native tunica media layers, thus the scaffolds produced herein closely match the structure of the native vessel wall. These architectures enabled an anisotropic mechanical behaviour to be obtained with PCL. Indeed, the major limitation of current available VG is the mismatch in the mechanical properties when compared to anisotropic behaviour of native vessels [4][10]. A smaller WA resulted in greater differences between circumferential and longitudinal moduli, simulating the contribution of structural proteins – collagen and elastin [10][55]. Interestingly, the anisotropic ratio between the second elastic region is similar to previous MEW scaffolds [22][23], but still divergent from the biological control. Nonetheless, the dynamic mechanical analysis showed that the use of rhombus-like pores replicate the characteristic J-shape curve of native tissue for a physiological relevant strain range of 10-15% [43][56][57][58]. Furthermore, the quantification of the unload to load ratio – identifying energy dissipation and therefore plastic deformation [59] – reported an increase from 5% to 15% strain in the circumferential direction, which occurred only beyond 35% strain for the longitudinal direction. Similar findings were reported also for wavy-like MEW PCL samples, mimicking heart valve tissue [22], whereby plastic deformation was reached at 15% and 20% for the two directions of testing. In summary, MEW technology demonstrated high accuracy in the fabrication of different scaffold architectures which provided an increased vasculature mimetic anisotropic mechanical response as the WA was reduced to match native collagen disposition.

ECM inspired MEW architectures can significantly enhance cell organisation, ECM deposition and orientation into highly organised structures. F-actin staining demonstrated that the chosen scaffold architectures can tune the cell spatial arrangement, in agreement with previous *in vitro* studies [28][40][51][53][60]. A square-shape pore was responsible for an orthogonally oriented distribution of the cell cytoskeleton [61][62], while the use of diamond-like pores allowed for a preferential cell organisation. The narrower the printed WA the higher the obtained alignment of cell along the major diagonal demonstrating the contact guidance provided by the MEW framework. Similar cell responses were reported in different diamond-shape MEW printed architectures [24][28][29], whereby the smallest angle reported and tested was 30° degrees [29]. Interestingly, histological staining highlighted that ECM was preferentially deposited parallel to that of the cell orientation. Importantly, polarised light microscopy validated that neo-tissue deposition was occurring circumferentially, as per the tunica media configuration. This native organisation of newly formed tissue is a good indicator of the longer-term survival of the tissue post graft degradation upon implantation. Interestingly, the WA-18° was able to induce ECM deposition towards the major diagonal of its architecture with a fluctuation around ± 15° degrees which corresponds very closely to native collagen organisation identified with diffusion tensor imaging-derived tractography [42][63]. Thus, demonstrating how the chosen graft, despite targeting 18° degrees, is specifically tuning collagen fibres deposition as per native collagen organisation *in vivo*. The best performing WA was identified in its ability to overcome the current limitation of SVGs, with enhanced mechanical properties and tissue regeneration capacity, hereby finding a valuable substitute for current therapeutical treatments.

VGs are tubular structures and therefore a translation from the optimal planar configuration to a tubular format was completed. SDVGs substitutes require diameters below 6 mm and an average length of 5 cm with maximum at 25 cm [9]. The tubular graft here printed with high accuracy satisfied this requirement with a 10 cm length and 3 mm internal diameter. Many groups have recently fabricated tubular constructs using MEW. Rod size typically spans between 16 mm [25] and 0.5 mm [28] in diameter; WAs are commonly between 70° [28] and 5° [64] and the typical height is around 10 stacked layers [25]. The variability of these grafts proves MEW to be a versatile platform for different uses in tissue regeneration (i.e., kidney tubule [28]). The current work is in line with such ranges of scaffold architectures but outperform the average in terms of pore size area – below the 0.5 mm^2^ reported [25] – and length. Remarkably, the results here presented showed a match in the stress-strain behaviour of native tissue until 10% strain and non-significant differences between native and polymeric grafts for the elastin dominant region, the physiological region, and ultimate tensile strength (UTS). It has been shown that different printing geometries affect the stress-strain response of tubular grafts [27], which gives a versatile platform for different engineered tissue. Together, the analysis of the mechanical behaviour here reported show that tuning scaffold geometry the J-shape response and the anisotropy of native vessels were matched. Once populated with cells, such structures will provide a suitable guidance for the deposition of ECM along the desired direction and recapitulate the mechanical response of native tissue.

## 6 Conclusion

MEW technology was adopted to create different architectures which represent the range of collagen orientations identified within the tunica media of blood vessels. These scaffolds demonstrated anisotropic mechanical behaviour resembling the mechanical response of native tissue for low levels of strain. Moreover, the adopted diamond shape drove cell infiltration and alignment along the major diagonal creating a boundary system where collagen fibre deposition occurred in the circumferential direction along the same angle present in the tunica media of the blood vessel wall. This MEW tunica media graft overcomes many of the current limitations of synthetic and electrospun TEVG, and thus may represent an important framework in the development of alternative small diameter vessel substitution grafts.

## Acknowledgements

This work was supported with by the Science Foundation Ireland (SFI) under [grant number 12/RC/2278 and 17/SP/4721] and co-funded by the European Regional Development Fund and SFI under Ireland’s European Structural and Investment Fund. This research has been co-funded by Johnson & Johnson 3D Printing Innovation & Customer Solutions, Johnson & Johnson Services Inc.

## Author contributions

ASF: design and fabrication of scaffolds, morphological analysis, SEM images mechanical and biological characterisation, tubular MEW fabrication and characterisation, data collection and analysis. BT, TL: MRI data collection and analysis. ASF, DAH, DJK: Conception and design of project flow. ASF: wrote the paper with input from all authors.

